# Contacting domains that segregate lipid from solute transporters in malaria parasites

**DOI:** 10.1101/863993

**Authors:** Matthias Garten, Josh R. Beck, Robyn Roth, Tatyana Tenkova-Heuser, John Heuser, Christopher K. E. Bleck, Daniel E. Goldberg, Joshua Zimmerberg

**Author notes:** co-Senior Authors (Daniel E. Goldberg, Joshua Zimmerberg).

## Abstract

While membrane contact sites (MCS) between intracellular organelles are abundant^1^, and cell-cell junctions are classically defined^2^, very little is known about the contacts between membranes that delimit extracellular junctions within cells, such as those of chloroplasts and intracellular parasites. The malaria parasite replicates within a unique organelle, the parasitophorous vacuole (PV) but the mechanism(s) are obscure by which the limiting membrane of the PV, the parasitophorous vacuolar membrane (PVM), collaborates with the parasite plasma membrane (PPM) to support the transport of proteins, lipids, nutrients, and metabolites between the cytoplasm of the parasite and the cytoplasm of the host erythrocyte (RBC). Here, we demonstrate the existence of multiple micrometer-sized regions of especially close apposition between the PVM and the PPM. To determine if these contact sites are involved in any sort of transport, we localized the PVM nutrient-permeable and protein export channel EXP2, as well as the PPM lipid transporter PfNCR1. We found that EXP2 is excluded from, but PfNCR1 is included within these regions of close apposition. Thus, these two different transport systems handling hydrophilic and hydrophobic substances, respectively, assume complementary and exclusive distributions. This new structural and molecular data assigns a functional significance to a macroscopic membrane domain.

## INTRODUCTION

Morbidity and mortality in malaria are due to the apicomplexan *Plasmodium spp.* replicating within the host RBC. During its initial invasion of the erythrocyte, the parasite invaginates the RBC plasma membrane to form the PV as a second barrier^3,4^. The parasite must install its own, unique transport systems in order to import and export everything it needs for its survival and proliferation^5–7^. Understanding the underlying transport mechanisms between the parasite and its host is useful to identify drug targets. Yet, these crucial transport systems remain incompletely understood for both hydrophilic and hydrophobic substances.

A PVM channel, permeable to water-soluble nutrients like monosaccharides and amino acids^5^, is formed by the ‘exported protein 2’ or EXP2^8^. EXP2 also facilitates protein export, by serving as the protein-permeant pore for the ‘*Plasmodium* translocon of exported proteins’ (PTEX)^9^. A number of PPM channels, including several that are specific for particular nutrients have been identified and studied^10,11^. However, it is not known how lipid substances are transported across the PV, since the two limiting membranes have never been seen to connect or to transport membrane vesicles between each other^6^. The PPM resident protein ‘*Plasmodium* Niemann-Pick C1-related protein’ (PfNCR1) is essential for lipid homeostasis^12^ but it is unknown how it functions. It is not clear how the PV is organized, in order to support the transport of such a large variety of substrates.

Here we hypothesize that the structure of the PV is built such that it can support direct exchange of lipids across the PV space, directly between the PPM and PVM, in regions that can be defined as membrane contact sites (MCS).

## RESULTS

### Regions of close PVM-PPM apposition exist

Thin-sections of parasitized red cells were examined in the electron microscope (EM) to determine the separation distance between PVM and PPM. Since transport across the PV is most active in the trophozoite stage, i.e. the stage when parasites have begun to accumulate hemozoin and grow the fastest but have not begun to divide^13^, only this stage was considered. An example image is shown in (Figure 1A). The distribution of separation distances between PVM and PPM in such organisms was found to be bimodal, indicating two distinct structural regions: regions of “close membrane apposition”, separated by ∼10 nm, and regions of “PV lumen” with wider membrane separation distances of 20-30 nm. To control for possible artifacts introduced by chemical fixation, infected red cells were also prepared by a quick-freeze, freeze-fracture method that preserved cells in their most lifelike state. Still, these two distinct types of regions could be clearly discerned (Figure 1B). In conclusion, these two distinct differences in separation are narrow enough that they could be bridged by protein complexes that could interconnect the PVM and PPM membranes, and thus are candidates for membrane contact sites^1^.

**Figure 1.**
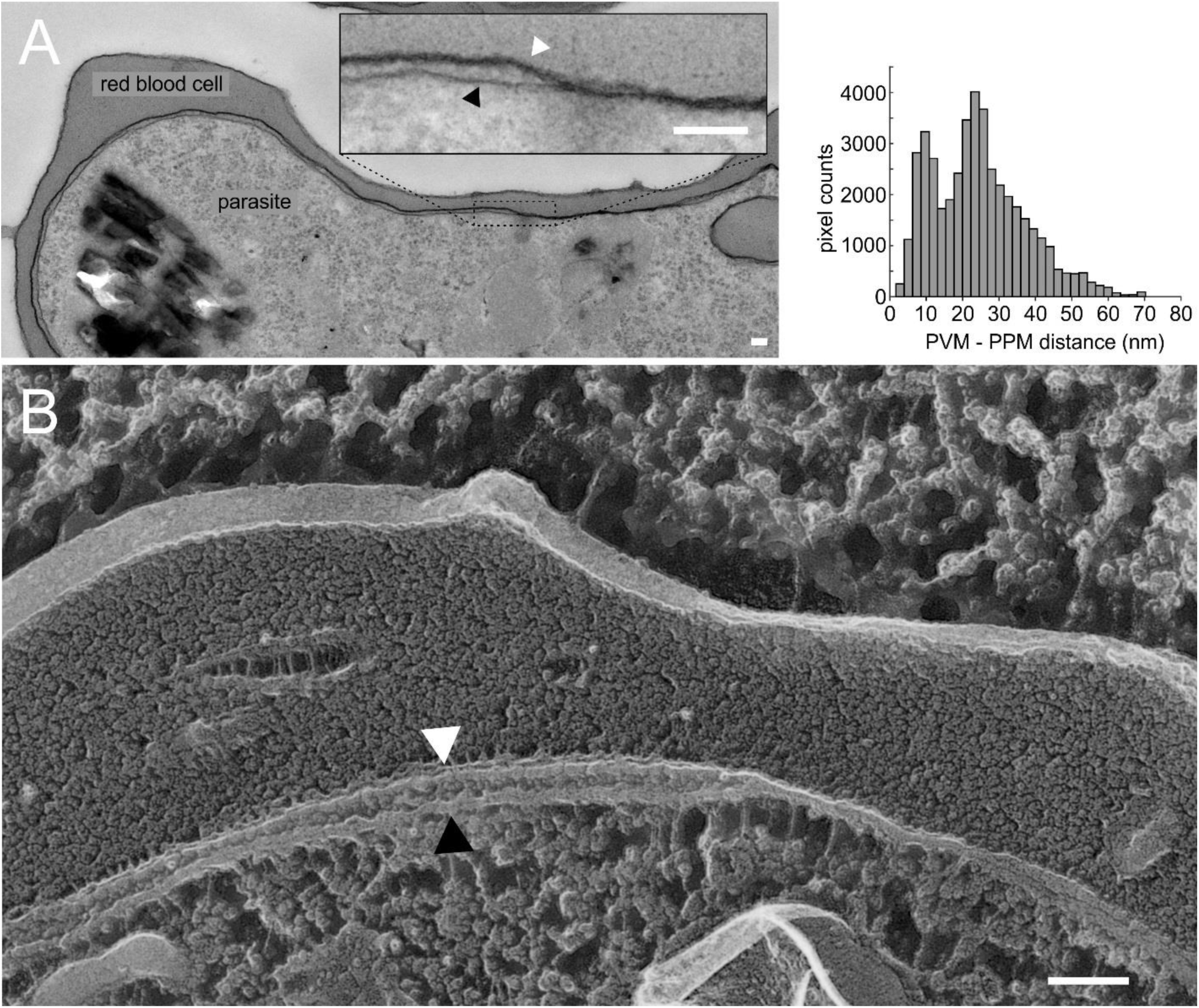
The PVM is wrapped tightly around the parasite. (A) (left) Thin section electron micrograph of a *Plasmodium falciparum* (NF54attb)-infected red blood cell. The inset highlights the PVM and PPM. (right) Histogram of the PVM-PPM distance collected from single sections of 7 parasites in 6 images in regions where both membranes are cut at a right angle. (B) Freeze fracture of NF54attb. A region of PV lumen is visible in the left of the image and a region of close membrane apposition is visible in the right side. (A and B) Scale bars: 100 nm. White arrowheads to the PVM, black arrowheads to the PPM.

### EXP2 localizes in domains of the PVM

A functionally characterized marker of the PVM, EXP2^8^, was used to investigate the domain structure of the PVM. To test domain formation in fixed but label-free, unmodified parasites (NF54), an immunofluorescence assay using EXP2 antibodies was performed. EXP2 was found in patches, interrupted by stretches of PVM devoid of anti-EXP2 label (Figure 2A). To test if these regions of EXP2 were domains in living parasites and to control for chemical fixation artefacts, a parasite line bearing a C-terminal mNeonGreen (mNG) fusion to the endogenous copy of EXP2 (EXP2-mNG) was examined^14^. In these parasites, the mNG signal also was found in continuous patches, interrupted by stretches of PVM devoid of mNG (Figure 2B). To quantify and visualize these apparent domains of EXP2, we developed a simple tool: a projection of the maximum fluorescence intensity from the inside of the parasite onto a sphere. This results in a map of the fluorescence signal of the periphery of the parasite (Figure 2 C-E). Both samples show a protein domain coverage on the order of 50% around the parasite (Figure 2F).

**Figure 2.**
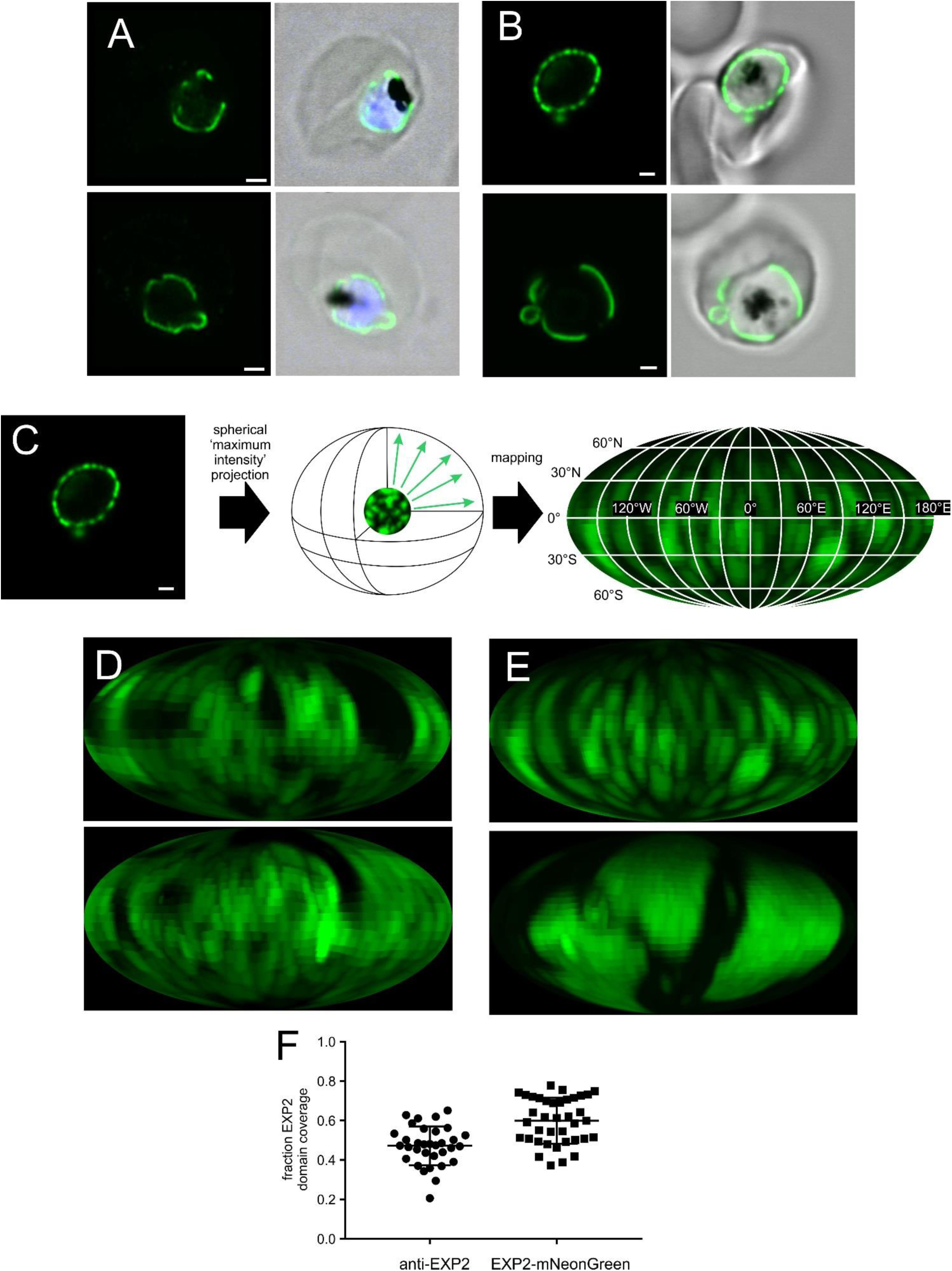
EXP2 is distributed in domains on the PVM. (A) Single confocal slices of an indirect immunofluorescence assay of NF54 showing EXP2 (green), DAPI (blue) and bright field (gray). (B) Single confocal slices of a live cell image of EXP2-mNeonGreen. EXP2-mNeonGreen (green), bright field (gray). (A&B) Scale bar: 1 µm. (C) Principle of the spherical intensity projection and mapping for the 2-dimensional analysis of the 3-dimensional dataset (see methods for details). (D&E) Mollweide projection of the images in A&B respectively. (F) Fraction of EXP2 domain coverage (mean+/-SD) in the projections determined by thresholding. The plot compares the domain coverage as measured by either NF54 antibody labeled against EXP2 (N=33 cells) and Exp2-mNeonGreen--PV-mRuby3 parasite line (N=38 cells).

Two physically independent techniques to visualize EXP2 spatial organization indicate its domain structure. Thus, EXP2-mNG can serve as a robust and readily detected marker for such PVM domains, labeling regions of protein export and transport of small water-soluble molecules.

### EXP2 co-localizes with PV lumen

The EM-findings of two regions with distinct PVM-PPM separation-distances and the light microscopy findings of domains for protein export and nutrient import represented by EXP2 led us to hypothesize that these features could be correlated.

To determine the distribution of the EXP2-domains relative to PV domains with distinct lumenal space, their localization was carefully mapped. Thus, a fluorescent label for the PV lumen was required that could be compared with the distribution of EXP2-mNG. To this end, mRuby3 was targeted to the PV lumen using the signal peptide of HSP101 (PV-mRuby3)^15^. To verify that PV-mRuby3 would serve as a genuine label of the PV lumen, correlative light/electron microscopy after cryo-thin sectioning was performed^16^. PV-mRuby3 was indeed found to localize to the regions of wider separation between PVM and PPM, confirming that it would serve a label for the accessible PV space (Figure 3A).

**Figure 3.**
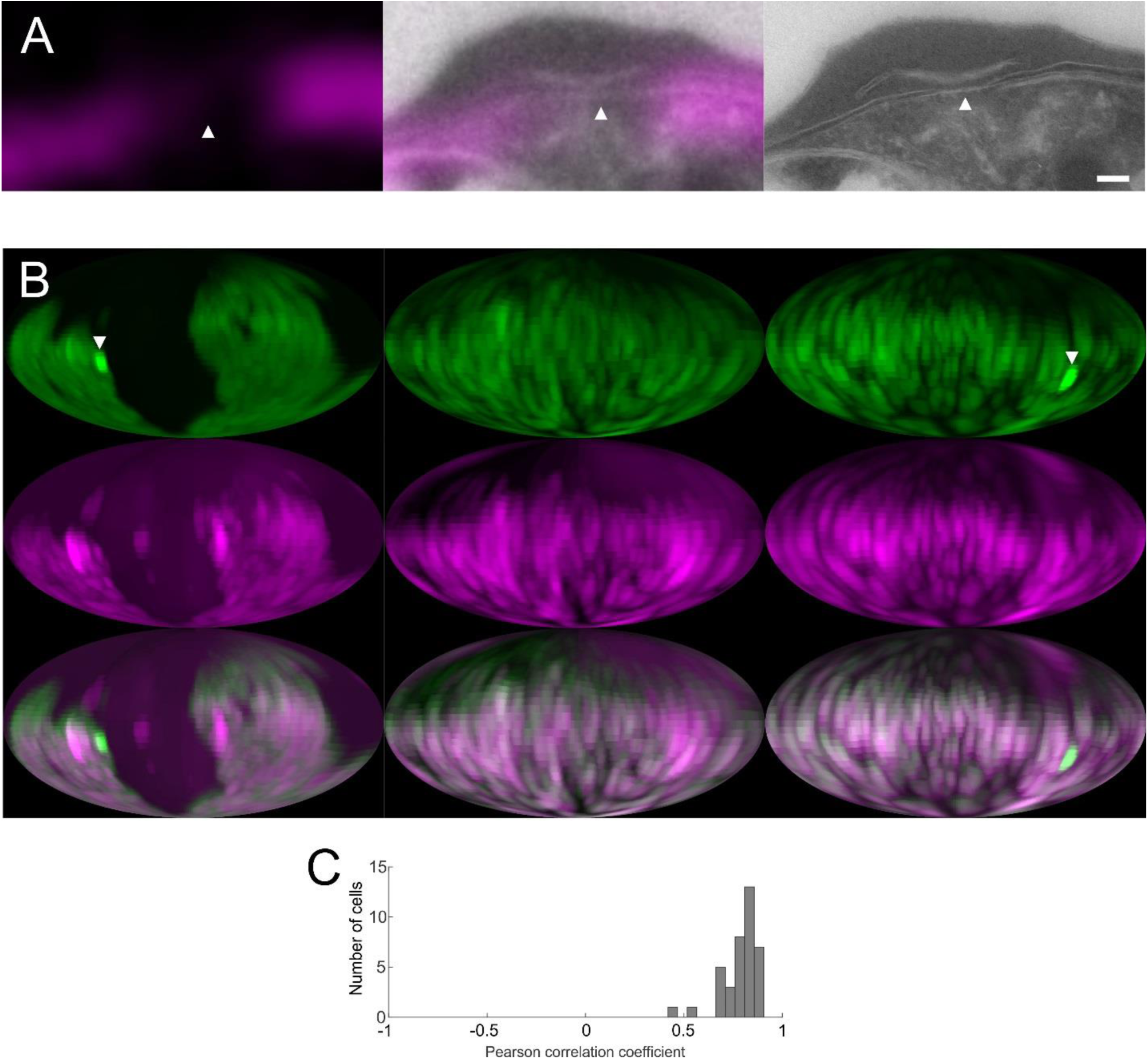
Colocalization of vacuolar space with EXP2. (A) Detail of correlative light electron microscopy of EXP2-mNeonGreen—PV-mRuby3 parasites showing the presence of PV-mRuby3 in the PV lumen and absence in the apposed membrane region (arrowhead) (see Figure SI 1 for the full image). (magenta, left) mRuby3 signal in the Tokuyasu cryo-section. (middle) Overlay of confocal and electron microscopy image. (right) Higher resolution electron microscopy image of the region. Scale bar: 100 nm. (B) Mollweide projections of Exp2-mNeonGreen—PV-mRuby3 parasites. Exp2-mNeonGreen (green, top), PV-mRuby3 (magenta, middle), merge (bottom). Samples chosen represent examples of relatively low (0.55), average (0.72) and high (0.85) Pearson correlation coefficients of the maps. White arrowheads show tubulovesicular network^44^ that has a relatively higher density of EXP2. (C) Histogram of Pearson correlation coefficients of trophozoite stage EXP2-mNeonGreen—PV-mRuby3 parasites (N = 38 cells). Regions that extend from the parasite to form the tubulovesicular network were excluded for the correlation analysis.

Co-expression and two-color imaging of EXP2-mNG and PV-mRuby3 demonstrated a clear-cut colocalization of both labels around the periphery of the parasite (Figure 3B). The mean Pearson-correlation coefficient for the analyzed sample is 0.81 CI [0.78, 0.83] (Figure 3C) as a mathematical measure for the degree of signal overlap, with numbers from −1 (perfectly anti-correlated), 0 (not correlated) to 1 (perfectly correlated).

Thus, EXP2 distribution can be correlated with regions of the wide mRuby3 accessible lumen exists. Within the limits of optical microscopy, they colocalize. Therefore, protein export, nutrient import, and presumably aqueous waste export occurs in the regions of the PV lumen.

### The lipid transporter PfNCR1 anti-localizes with EXP2 and the PV lumen

Sites of close membrane contact are implicated in direct transfer of lipids via intervening or included proteins that localize to those domains^1^. In the PPM PfNCR1 has been found to be essential for the maintenance of lipid homeostasis^12^. However, its human homolog, Niemann-Pick C1 (hNPC1) relies on a co-factor hNPC2, to transport lipids^17^; no such co-factor has been identified in *Plasmodium spp.*, suggesting that PfNCR1 may work differently. Thus, localizing PfNCR1 with respect to the separation-distances of the membranes of the PV is indicative in order to determine if PfNCR1 can be expected to transport lipids by interacting with a soluble cofactor from the PV lumen, or whether it interacts directly with the PVM at sites of close membrane apposition.

To determine the distribution of PfNCR1 relative to the domains defined by EXP2 and the PV lumen, an endogenous EXP2-mRuby3 fusion was engineered into a parasite expressing an endogenous PfNCR1-GFP fusion protein, allowing both proteins to be monitored by live fluorescence while preserving their native timing and expression levels. Two-color imaging showed that localization of the two labels is anti-correlated around the periphery of the parasite (Figure 4A). The mean Pearson-correlation coefficient of this sample is −0.20 CI [−0.10, −0.29], giving a mathematical measure for the anti-correlation of both signals (Figure 4B). Positive values of the coefficient are caused by small domains, approaching the resolution limit of light microscopy with signals co-localizing at the border of domains (Figure 4A). Anti-correlation was also observed in parasites where the PV-mRuby3 lumenal reporter was expressed in the PfNCR1-GFP background (Figure SI 2). From this observed anti-localization of the lumenal *vs.* hydrophobic transporter labels, it is most likely that PfNCR1 is localized to regions of close membrane apposition.

**Figure 4.**
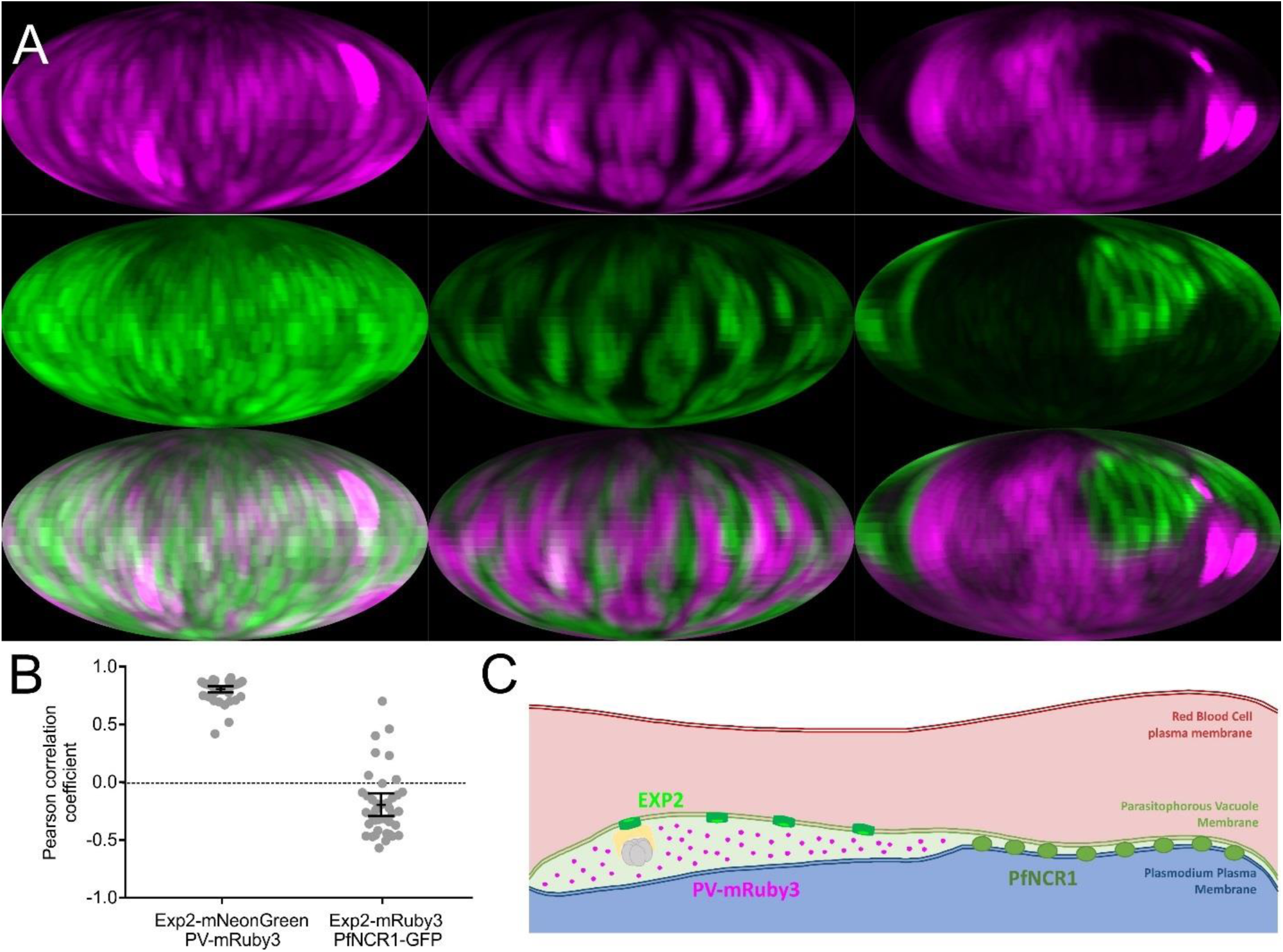
Anti-colocalization of EXP2 and PfNCR1. (A) Mollweide projections of PfNCR1-GFP—EXP2-mRuby3 parasites. EXP2-mRuby3 (top, magenta), PfNCR1-GFP (middle, green), merge (bottom). Samples chosen represent examples of relatively low (0.46), average (−0.22) and high (−0.71) anti-correlation according to the Pearson correlation coefficients of the maps. (B) Pearson correlation coefficients of Exp2-mNeonGreen—PV-mRuby3 parasites (N = 38 cells, see figure 3) in comparison to PfNCR1-GFP—EXP2-mRuby3 (N=39 cells). Regions that extend from the parasite to form the tubulovesicular network were excluded for the correlation analysis. Bars: mean +/- 95% CI. (C) Localization of EXP2 correlates with the existence of a relatively large PV lumen. The PV lumen can store proteins for export and accessory proteins to facilitate protein export. The lipid transporter PfNCR1 localizes to regions of close PPM-PVM apposition. Functionally, PfNCR1 may be able to directly contact the PVM to exchange lipids and sites of membrane apposition may be sites for the general exchange of lipophilic material, a functional hallmark of membrane contact sites.

## DISCUSSION

In studying how the malaria parasite modifies its protective membrane barriers to import and export materials that it needs to survive and grow, we found that the host cell-parasite interface (HPI) is a unique intercellular junction with clearly definable regions of protein composition and variable separation, consisting of a parasite vacuolar membrane, the lumen of the vacuole (that is the parasite’s extracellular space), and the plasma membrane of the parasite. Different classes of molecules, hydrophobic and hydrophilic, are transported through this one continuous, spheroidal interface with distinct regions. This segregation of function is accomplished by the creation of membrane contact sites (MCS) like those between cellular organelles^18,19^. Electron microscopy resolved the PVM and the PPM, allowing quantification of their separation-distances, and confirming the visual impression that the PVM and the PPM form distinct domains, characterized by a bimodal distribution of space between the two membranes. Using two complimentary techniques, indirect immunofluorescence and live-cell microscopy of fluorescently tagged EXP2, this solute-transporter was detected in µm-sized domains that correlate spatially with domains where the PVM and the PPM are separated enough from each other for the vacuole to accumulate a visible amount of the lumenal marker PV-mRuby3. In contrast, we found that the parasite lipid transporter PfNCR1 was specifically excluded from these regions and instead accumulated in the intervening regions of close PVM-PPM apposition. We conclude that the PV has evolved to become laterally segregated into relatively wide regions for hydrophilic transport, and separate close-contact regions for hydrophobic transport.

The distances between the PPM and PVM for both domains could be bridged by proteins, potentially qualifying both regions as MCS^1^. However, in neither domain of the HPI were bridging proteins observed in our “deep-etch” EMs, nor were they observed in our thin-section EMs. However, when the PV is induced to swell experimentally, e.g., when PVM protein export is conditionally impaired, leading to protein accumulation^8,20,21^, it expands inhomogenously into irregular protuberances, suggesting that some sort of adhesion normally exists between the PPM and the PVM that prevents it from swelling uniformly. Still to be determined is whether these adhesions concentrate in the tightly apposed regions or the more open regions. EXP2-mNG is included in the distended regions of the PVM when protein export is impaired (Figure SI 3), indicating that the EXP2-containing domains form the less strongly connected region.

In contrast, and in keeping with the detrimental effect of water on lipid transport, sites of close PVM-PPM apposition seem to be devoid of PV lumen altogether. This is an extremely close apposition, at the lower end of the membrane distances found at organelle-organelle MSC, i.e. 10-80 nm^1^. PfNCR1, a lipid transporter, localizes to these sites of unusually close membrane apposition. While it can be inferred that lipid transport is taking place at these sites, it is unclear how PfNCR1 goes about exchanging its substrate, presumably cholesterol, with the PVM. It has been demonstrated that the large extra-membranous domain of PfNCR1 is localized in between PPM and PVM. The closely apposed membranes are within the proteins hypothesized radius^12^, so it may well hand over lipids directly^22^, or exchange lipids with the help of a membrane-bound cofactor. PfNCR1 may also function as an anchor between these two membranes, given the relatively constant separation between them at the close contacts. Curiously, the domain structure of the PVM exemplified by the EXP2 distribution shown here, is quite dynamic and variable (*cf.*, SI movie, demonstrating remarkable flexibility in the PV), suggesting there may exist an active mechanism of protein localization driven by processes in the parasite cytoplasm. Additionally, proteins may target to their respective regions by various other mechanisms, such as interacting with structural proteins, sensing of membrane distance, as found with other contact sites^1^, or protein exclusion, as found at gap junctions^2^.

The structure-function relationship described here (Figure 4C) can potentially guide the study of other functions at the HPI, such as the inhomogeneously distributed PVM proteins noted previously^23,24^. A mechanism for the transport of resident membrane proteins from the PPM to the PVM is still lacking as PTEX is not involved in this process^25,26^. The regions of close apposition are a promising place to look for machinery that would allow transfer of such proteins.

The size of these domains are similar to those created in lipid phase demixing seen in model membranes^27^. However, cellular membranes are complex mixtures that for the most part fail to demix at physiological temperatures and remain almost entirely liquid-disordered^28^. Moreover, cholesterol-rich domains are not detected by surface imaging using mass spectroscopy^29^. However, molecular dynamic simulations have revealed nanoscopic microdomains of hexagonally packed saturated lipids^30^ that may correspond to the nonideal demixing revealed by FRET^31^. Recently a fungal vacuole was reported to exhibit temperature-dependent lipid phase demixing^32^ but no function has yet been ascribed to these domains. The specific role of lipid asymmetry and composition, and its role in domain formation and maintenance, in the face of lipid transport, remain to be investigated for any MCS or membrane domain. The HPI offers a larger platform for these studies that may benefit both cell biological and medical investigations.

## Supporting information

Figure SI movie. Dynamics of relatively small EXP2 domains.

## ACKNOWLEDGEMENTS

We thank Svetlana Glushakova for invaluable help with the cell culture and many discussions in the lab and Paul Blank for help with statistical questions. This work was supported by the Division of Intramural Research of the *Eunice Kennedy Shriver* National Institute of Child Health and Human Development, National Institutes of Health and National Institutes of Health grant HL133453 to J.R.B.

## CONTRIBUTIONS

M.G., J.Z., D.E.G. and J.R.B. conceived and designed experiments. J.R.B. generated and analyzed the parasite strains, performed light microscopy of the DHFR strain, generated the samples for the freeze fracture. M.G. performed all other light microscopy, the IFA, analyzed all microscopy data. R.R. prepared the freeze-fractures. T.T.-H. embedded and cut the thin-sections for EM, J.H. imaged freeze-fractures and thin sections. C.K.E.B. processed the sample for CLEM, imaged the sample in EM. All authors discussed and edited the manuscript.

## CODE AVAILABILITY

MATLAB (Mathworks) scripts as described in the Methods are available from the corresponding authors on request.

## DATA AVAILABILITY

The data that support the findings of this study are available from the corresponding authors on request.

## METHODS

### Cell culture

Plasmodium falciparum was cultured in T25 Nunclon Delta closed cap culture flasks (Thermo Fisher, Waltham, MA) in RPMI 1640 supplemented with 25 mM HEPES, 0.1 mM hypoxanthine, 25 μg/ml gentamicin (all Thermo Fisher), 0.5% Albumax II (Gibco, Gaithersberg, MD), and 4.5 mg/ml glucose (MilliporeSigma, St. Louis, MO) at 37 °C in a 5% CO2, 5% O2 atmosphere at 5% hematocrit below 5% parasitemia. Red blood cells were obtained from anonymized healthy donors in a protocol approved by the National Institutes of Health Institutional Review Board. NF54 obtained through BEI Resources, NIAID, NIH as part of the Human Microbiome Project. The parasite lines Exp2-mNeonGreen^14^, Exp2-mNeonGreen—PV-mRuby3^15^, NF54attb^33^ were described previously.

### Molecular Biology

For generation of an endogenous EXP2-mRuby3 fusion, the blasticidin-S deaminase (BSD) cassette between SalI and BglII in the plasmid pGDB^34^ was isolated by digestion and gel purification and inserted between the same restriction sites in the plasmid pyPM2GT-EXP2-mNeonGreen^14^ with a T4 Quick Ligation kit (NEB), replacing the yDHODH cassette. A synonymous mutation in the EagI site within the BSD coding sequence was then introduced to inactivate this site using a QuikChange Lightning Multi Site Directed Mutagenesis kit (Agilent, Santa Clara, CA) and the primer GCCAGCGCAGCTCTCTCTAGCGACGGGCGCATCTTCACTGGTGTCAATG. The mRuby3 coding sequence was then PCR amplified from plasmid pLN-HSP101-SP-mRuby3^15^ with primers GATGAAAATAAAGAACCTAGGGGAAGTGGAGGAGTG and TAACTCGACGCGGCCGTCACTTGTACAGCTCGTCCATGCC and inserted between AvrII and EagI using an In-Fusion cloning kit (clontech), replacing the mNeonGreen coding sequence and resulting in the plasmid pbPM2GT-EXP2-mRuby3. This plasmid was linearized at AflII and co-transfected with the pAIO-EXP2-CT-gRNA^14^ into a parasite line bearing a C-terminal GFP fusion to the endogenous *pfncr1* gene^12^ and selection with 2.5 µg/ml blasticidin-S was applied 24 hours after transfection. For expression of PV-targeted mRuby3, the plasmid pLN-HSP101-SP-mRuby3 was co-transfected with pINT^35^ into the PfNCR1-GFP background and selection with 2.5 µg/ml blasticidin-S was applied 24 hours after transfection.

To generate an endogenous EXP2-mNeonGreen fusion in the HSP101^DDD^ background, the plasmid pyPM2GT-EXP2-mNeonGreen was co-transfected with pAIO-EXP2-CT-gRNA into NF54attB: HSP101^DDD 8^. Selection was applied with 2µM DSM1 24 hours post transfection and parasites were cloned by limiting dilution when they returned from selection.

### Immunofluorescence assay (IFA)

IFAs were performed as described previously^8^. Briefly, cells taken directly from the culture and left to settle for 10 min on Concanavalin A (Vector Laboratories, Burlingame, CA) coated cover slides in culture medium at 37°C. Excess cells were taken off with 3 gentle washes using 37°C cell culture medium. Cells were fixed for 15 min at 37°C in freshly prepared 4% paraformaldehyde, 0.0075% glutaraldehyde (both from electron microscopy sciences), in phosphate buffered saline (PBS) (Gibco). After 3 washes in PBS cells were permeabilized in freshly prepared PBS containing 0.2% Triton X-100 (MilliporeSigma). The sample was then blocked in PBS + 3% Bovine Serum Albumin (BSA) (MilliporeSigma). Cells were incubated in primary Monoclonal antibody 7.7 (anti-EXP-2), obtained from The European Malaria Reagent Repository (http://www.malariaresearch.eu, 1:500) for 1 hr at room temperature. The sample was washed 5 times in PBS. The secondary antibody (Donkey-anti-mouse conjugated with Alexa Fluor 488, from Invitrogen, lot 1113537, 1:150) was applied for 20 min, then washed off 5 times with PBS. The sample was mounted in ProLong AntiFade Gold with DAPI (Invitrogen).

### Light microscopy

Images were obtained on a Zeiss 880 with Airyscan module using a 63x 1.4NA Zeiss Plan-Apochromat, 37 °C immersion oil (Zeiss, Oberkochen, DE). Images were collected using Zen black (Zeiss) in the Airyscan mode, following the programs recommendation for the optimal pixel size and slice thickness, pixel dwell times were kept at 1-2 µs. Live parasites were transferred into a hybridization chamber (HybriWell HBW20, Grace Bio-Labs, Bend, OR) for observation on the microscope^15^. The stage was heated to physiological temperature using a stage incubator (Tokai Hit INU, Fujinomiya-shi, Japan) (set temperatures: top 37 °C, stage 36 °C, objective 39 °C) to reach 35-36 °C at the coverslip. Trophozoites, i.e. non-segmented cells with hemozoin, were chosen from a bright field image. For colocalization imaging following parameters were chosen: Imaging on EXP2-mNeonGreen—PV-mRuby lines was done using a “band pass 495-550 nm+ long pass 570 nm” emission filter, switching excitation at each line between 488 nm at 0.1% and 561 nm at 1% power. For experiments with GFP as fluorescent tag the line switch strategy, while minimizing movement artefacts, lead to blead through from the mRuby3 to the GFP channel as the mRuby3 signal was very abundant and mRuby3 can be minimally excited with 488 nm making it visible alongside the relatively weak GFP signal in the GFP channel. This required switching filters after completing z-stacks in each individual channel. Parameters chosen to image PfNCR1-GFP—EXP2-mRuby3 are for the GFP channel excitation: 488 nm at 2% power, emission: “band pass 420-488 nm + band pass 495-550 nm”, and the mRuby3 channel excitation: 561 nm 0.2%, emission: bandpass “495-550 nm+ long pass 570 nm”. The images were processed in Zen black using Airyscan processing in automatic settings.

### Map projection and correlation analysis

PVM, PV lumen and PPM are not resolvable from each other with a light microscope but appear as a single surface. To avoid thresholding for the correlation analysis of the light microscopy data, the 3-dimensional dataset was reduced to 2 dimensions projecting the maximum fluorescence intensity from the calculated center of the parasite onto an “infinite” sphere in 1° intervals in all directions. This information can then be used to draw an angular map of the signal. A Mollweide projection was chosen as it represents an equal-area projection when assuming a spherical parasite. Maps obtained this way are similar to how the night sky can be represented in a map^36^. The Pearson correlation coefficient was then calculated on the maps. The maps also provide an informative visual impression of the 3D dataset in print.

Analysis was done using custom scripts in MATLAB 8.5 (MathWorks, Natick MA). Briefly, the center of the cell was determined from the center of mass of a mask created from pixels in between the 1^st^ and the 2^nd^ level of a 2-level threshold using the “multithresh” function. Each voxel was then assigned an altitude and azimuth with respect to this center using the “acos” and “atan2” functions respectively. The altitude and azimuth were then used to create a mask with a 1° resolution (equivalent to 17.5 nm at a distance of 1 µm from the center). The maximum of masked intensity was then recorded as angular intensity value for the altitude and azimuth. Finally, the angular intensity values were mapped using a Mollweide projection onto a 1024×2048 pixel sized image. For each pixel with coordinates xi and yi corresponding angles were calculated:

~~~
ps = 1024;
R = ps/(2*sqrt(2));

y = ((yi-(ps/2))/(ps/2))*(R*sqrt(2));
th = asin(y/(R*sqrt(2)));
Latitude(yi,:) = real(asin((2* altitude +sin(2* altitude))/pi)*180/pi);
~~~

and

~~~
x = ((xi-ps)/ps)*(2*R*sqrt(2));
L = real((pi*x)/(2*R*sqrt(2)*cos(th))*180/pi);
Longitude(yi,xi) = real((pi*x)/(2*R*sqrt(2)*cos(th))*180/pi);
~~~

Area coverage of the signal was determined by segmenting the individual maps with the “multithresh” function for 2 levels, counting everything above the first threshold as signal. Correlation coefficients (r) were correlated using the Pearson correlation coefficient from the intensities of channel 1 (XL) and channel 2 (YL):

~~~
N = length(XL);
mX = mean(XL);
mY = mean(YL);
r = (sum(XL.*YL)-N*mX*mY)/(sqrt(sum(XL.*XL)-N*mX*mX)*sqrt(sum(YL.*YL)-
N*mY*mY));
~~~

Regions showing the tubulovesicular network, identified as PVM extending out from the parasite into the RBC cytosol, have enriched EXP2 signal compared to the peripheral PV and were manually masked out when calculating the correlation coefficient.

The 95% confidence interval of the correlation coefficient was calculated in Prism (Graphpad, San Diego, CA) after a Fisher transform of the data, the result was then back transformed. The source code of the scripts used will be made available through the authors and online repository.

### Electron Microscopy on thin sections

Erythrocyte cultures infected with NF54attb parasites were enriched for late stages using a LD column in a QuadroMACs magnetic separator (Miltenyi Biotech, Cologne, DE), then fixed with 2% glutaraldehyde in 100mM NaCl, 30 mM Hepes buffer pH 7.2, and 2 mM CaCl_2_. During a 1-2 day long aldehyde fixation, the cultures settled into soft pellets, after which they were post-fixed for 30 minutes as pellets in 0.25% OsO_4_ plus 0.25% potassium ferrocyanide dissolved in 0.1 M cacodylate buffer containing 2 mM CaCl_2_. Thereafter, they were washed in 0.1 M cacodylate buffer+2mM CaCl_2_, and poststained by sequential 30 min treatments with 0.5% tannic acid in 0.1 M cacodylate buffer+2 mM CaCl_2_ followed by 0.5% uranyl acetate in 50 mM acetate buffer pH 5.2. Immediately thereafter, they were dehydrated with increasing concentrations of ethanol, and embedded in epoxy resin by standard techniques: exchange through propylene oxide, then increasing concentrations of the epoxy, and final vacuum-embedding and polymerization at 70 °C. Thereafter, semi-thin sections were made vertically through the pellets at 0.5 µm and stained with toluidine blue, to determine optimal areas for further examination. These were then sectioned at 80 nm, stained for 5 min with 50 mM lead citrate dissolved in 0.1 M citrate buffer pH 5.2, and examined at 80 KV in a JEOL 1400 electron microscope equipped with a 4Kx6K digital camera (Advanced Microscopy Techniques Corp, Woburn). In digital versions of the final electron micrographs, PPM and PVM were segmented by hand, using ImageJ^37^, drawing a line where each membrane was clearly visible. Sections where the membranes were cut obliquely were not considered. The minimal distance of the two membrane traces was determined using a MATLAB script.

### Correlative Light Electron Microscopy

EXP2-mNeonGreen—PV-mRuby3 parasites were isolated on a 65% percoll (MilliporeSigma) interface. Cells were fixed using 4 % formaldehyde + 0.4 % glutaraldehyde in 1x PHEM (pH 6.9) (Electron microscopy sciences)^38^. Gelatin-embedded samples were infiltrated with 2.3 M sucrose (in 0.1 M phosphate buffer) and put at 4 °C for three days on a rotating wheel, mounted onto sample pins and frozen in liquid nitrogen. Subsequently, the samples were ultrathin cryo-sectioned (50–60 nm) with a FC7/UC7-ultramicrotome (Leica, Vienna, AT) and with a 35° diamond knife (Diatome, Hatfield PA), picked-up with a 1:1 mixture of 2% methylcellulose (25 centipoises, MilliporeSigma) and 2.3 M sucrose (USB Corporation, Cleveland, OH)^39^. Sections were 5-10 min washed in PBS prior to light microscopy imaging using the Zeiss LSM 880 Airyscan module (see the section on light microscopy). Samples were then embedded in 4% uranyl acetate/2% methylcellulose mixture (ratio 1:9)^40^ for electron microscopy. Thin sections were examined on a JEM-1200EX (JEOL USA) transmission electron microscope (accelerating voltage 80 KeV) equipped with an A.M.T. 6-megapixel digital camera (Advanced Microscopy Techniques, Woburn, MA). EC-CLEM^41^ was used to align light and electron microscopy image in a non-rigid grid to accommodate sample warping. Electron micrographs with up to 2500x direct magnification (7.4 nm pixel size) were used for the alignment to the light microscopy image (pixel size following the “optimal” pixel size settings of the Airyscan module is 42.6 nm).

### Freeze Fracture Replica

Late stage NF54attb infected red blood cells were isolated using a magnetic separator as for the thin sections. Cell were gently pelleted at 1000 RPM in a clinical centrifuge, then layered as a thick slurry on a tiny 3×3mm class coverslip mounted on a lung cushion, in preparation for quick-freezing with the liquid helium cooled copper-block “slammer”^42^. Thereafter, they were transferred to a Balzers 400 freeze-fracture apparatus, where they were fractured through their well-frozen surfaces, “deep-etched” for 2 min at −104 °C, and then rotary-replicated with 4 nm of platinum deposited from a 20° angle. Thereafter they were “backed” with 10 nm of carbon deposited from 90°, removed from the Balzers, thawed, and the platinum replica was floated on 25% SDS to partially remove organic material from underneath it^43^. After washing, the replica was picked up on an EM grid. Thereafter, these replicas were examined in the electron microscope in the same manner as the thin sections, above.

## FIGURE LEGENDS

**Figure SI 1.**
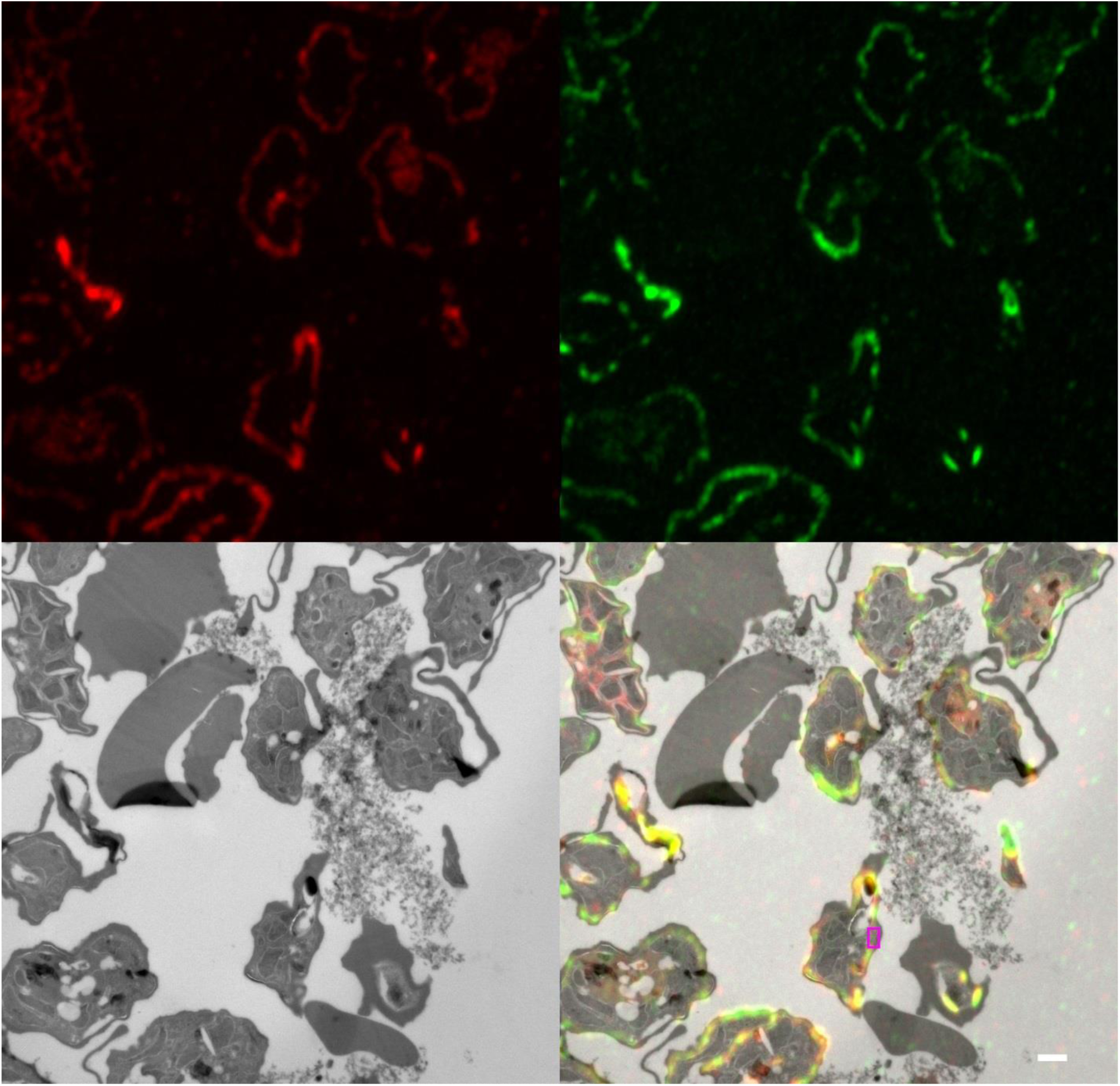
Correlative light electron microscopy. Zoomed out view of the correlation shown in figure 3. Red: PV-mRuby3, Green: EXP2-mNeonGreen, Gray: electron micrograph. The magenta box indicates the region shown in figure 3. Scale bar: 1 µm.

**Figure SI 2.**
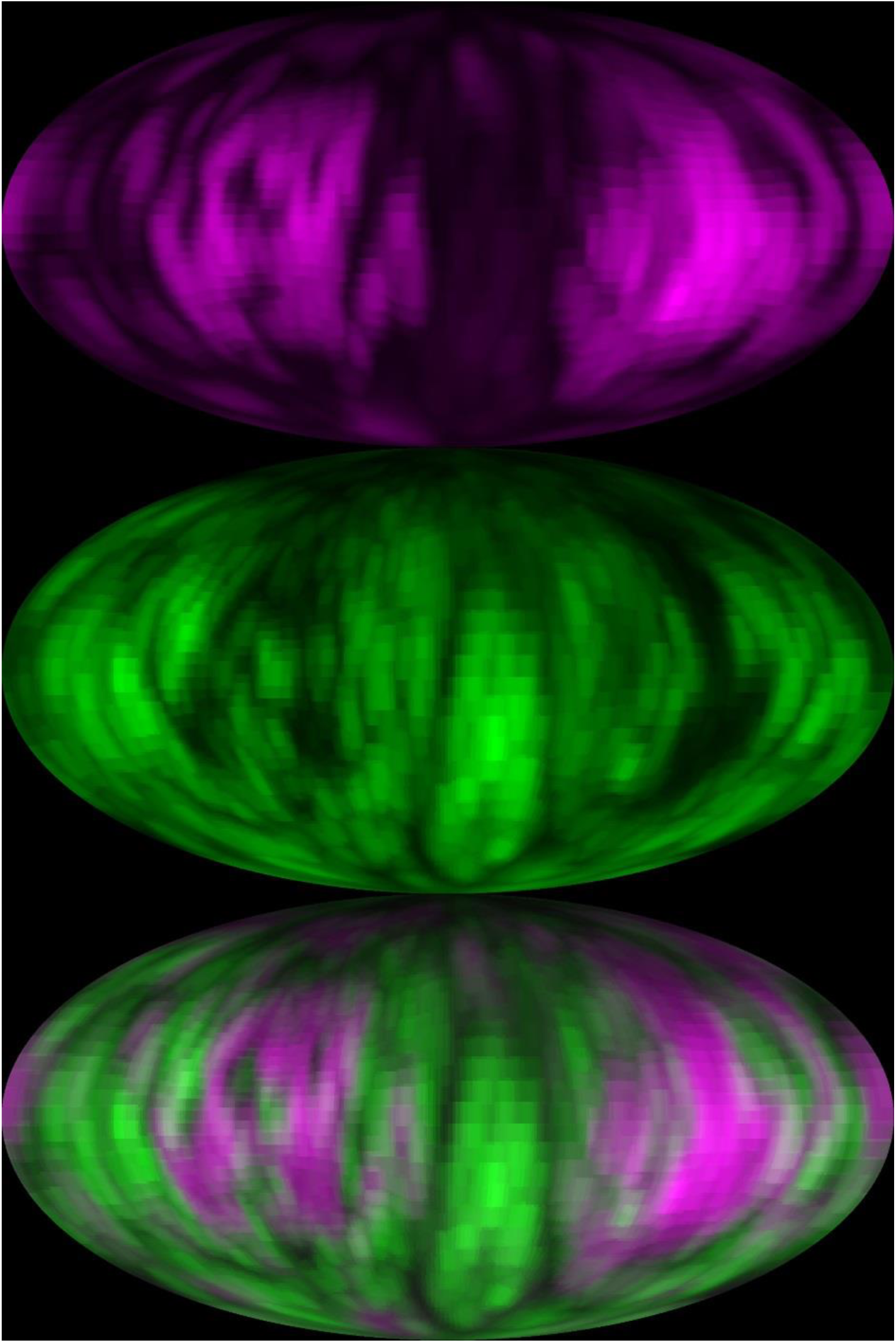
Correlation of PV-mRuby3 and PfNCR1-GFP. Mollweide projections of a parasite expressing PV-mRuby3 (magenta) and PfNCR1-GFP (green). The Pearson correlation coefficient for this cell is −0.20.

**Figure SI 3.**
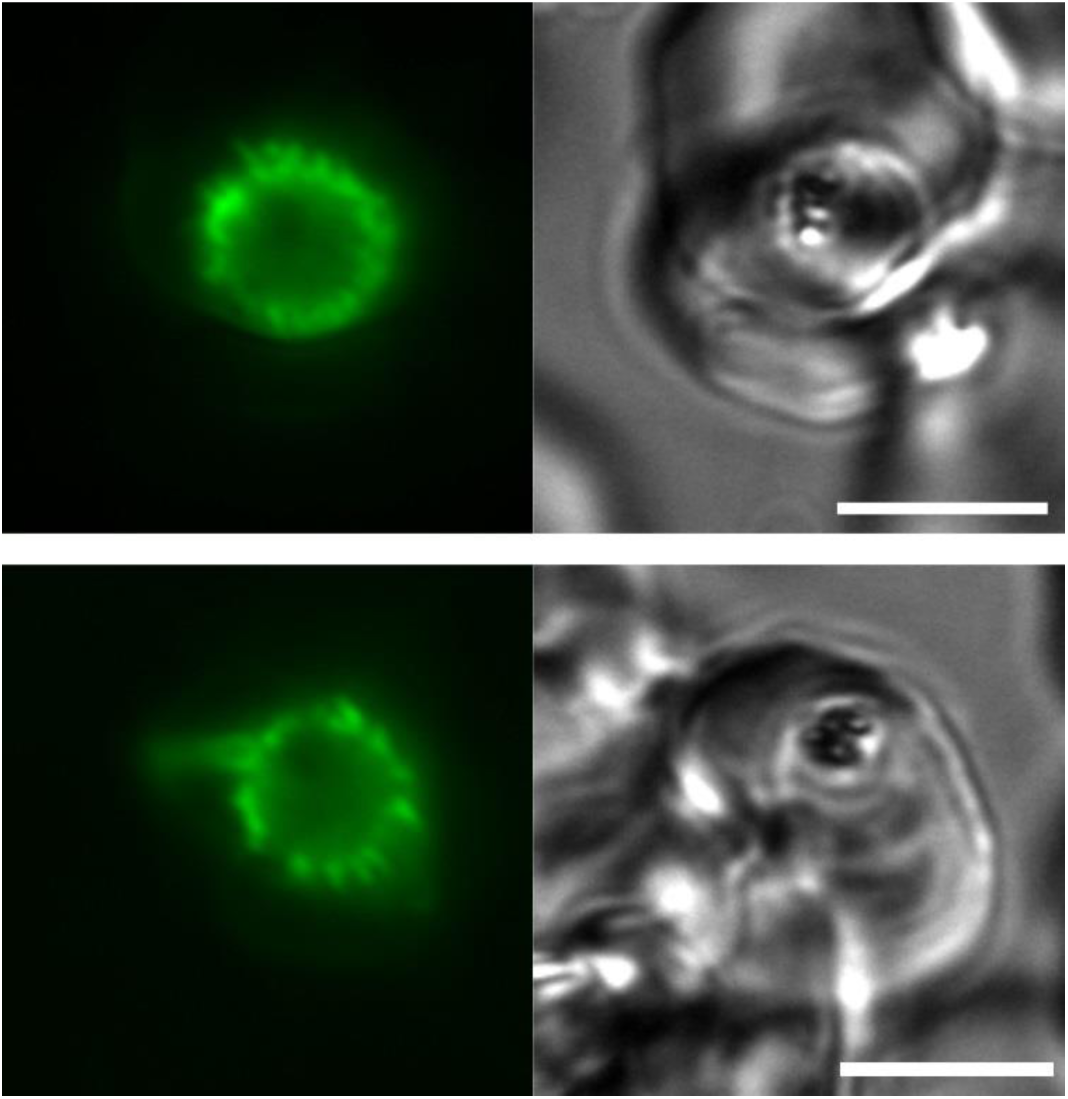
EXP2 is targeted to distensions when protein export is impaired. Compared to the broken but smooth outline of EXP2-mNeonGreen in unmodified parasites (see figure 2) the outline after impairing protein export by inactivating HSP101 using a (DHFR)-based destabilization domain (DDD) leads to the appearance of a distended signal. This suggests that EXP2 is present in the distended membrane reported in ^8^. Green: Exp2-mNeonGreen, gray: brightfield, scale bar: 5 µm. Trimethoprim was withdrawn 24 hours before imaging. Image was acquired on an Axio Imager M1 equipped with a Plan-Apochomat 100x/1.4NA objective and ORCA-ER CCD camera, using the GFP filterset for the mNeonGreen imaging.

**Figure SI movie. Dynamics of relatively small EXP2 domains.**

Time course of Z-projection of a stack of the PVM face closest to the cover slide. Green: EXP-mNG. Scale bar: 1 µm.

