## Supplementary figures and images for "Contacting domains that segregate lipid from solute transporters in malaria parasites"

### Figure SI movie. Dynamics of relatively small EXP2 domains.

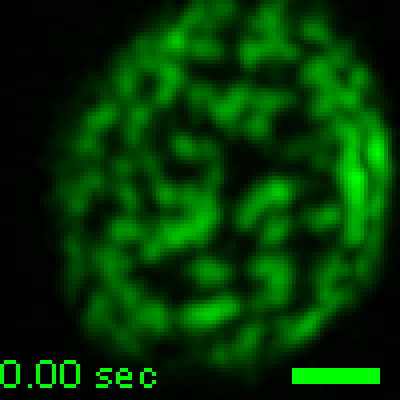
